# Enhanced MATE transporter DTX6/PQT15 confers paraquat resistance

**DOI:** 10.1101/2021.02.07.430154

**Authors:** Jin-Qiu Xia, Tahmina Nazish, Ayesha Javaid, Mohsin Ali, Qian-Qian Liu, Liang Wang, Zheng-Yi Zhang, Zi-Sheng Zhang, Yi-Jie Huang, Jie Wu, Zhi-Sen Yang, Lin-Feng Sun, Yu-Xing Chen, Cheng-Bin Xiang

## Abstract

Paraquat is one of the most widely used non-selective herbicides and has elicited the emergence of paraquat resistant weeds. However, the molecular mechanisms of paraquat resistance are not completely understood. Here we report an Arabidopsis gain-of-function mutant *pqt15-D* with significantly enhanced resistance to paraquat and the corresponding *PQT15* encoding the MATE transporter DTX6. A point mutation at +932 bp in *DTX6* causing the G311E amino acid residue change brings about the enhanced paraquat resistance of *pqt15-D*. Overexpression of *DTX6/PQT15* in the wild type also confers strong paraquat resistance, whereas the *DTX6/PQT15* knockout mutants exhibits hypersensitive phenotype to paraquat. Moreover, heterologous expression of *DTX6* and *DTX6-D* in *E. coli* significantly enhances bacterial resistance to paraquat. DTX6/PQT15 mainly localizes in the plasma membrane as shown by DTX6-GFP and functions as a paraquat efflux transporter as demonstrated by paraquat efflux assays with isolated protoplasts and bacterial cells. Taken together, our results demonstrate that DTX6/PQT15 is an efflux transporter and confers paraquat resistance by exporting paraquat out of cytosol, therefore unraveling a molecular mechanism of paraquat resistance in higher plants and providing a promising candidate of generating paraquat resistance crops.

## INTRODUCTION

Paraquat (or methyl viologen; 1,1′-dimethyl-4,4′ bipyridinium dichloride), a strong inducer of oxidative stress, is a widely used broad spectrum, quick-acting, nonselective herbicide. It quickly competes for electrons from PS I (photosynthesis system I) and reacts with oxygen, producing large quantity of reactive oxygen species (ROS) that destroy the photosynthesis system and ultimately kill plants (Hawkes, 2014; Suntres., 2002).

Paraquat resistance has been reported in weeds and Arabidopsis (Chen et al., 2009; Fujita et al., 2012; Itoh and Miyahara, 1984; Li et al., 2013; Luo et al., 2016; Murgia et al., 2004; Xi et al., 2012). Based on physiological studies of paraquat resistant weeds, several resistance mechanisms have been proposed, including translocation, sequestration, catabolism, and ROS scavenging (Fujita and Shinozaki, 2014; Hawkes, 2014; Murgia et al., 2004; Szigeti and Lehoczki, 2003). Although no genetic loci of paraquat resistance have been identified in weeds, some of the proposed mechanisms have been confirmed in Arabidopsis. For instance, *AtPDR11* encodes a plasma membrane-localized ATP-binding cassette transporter that uptakes paraquat from environment into cells. Therefore, knockout of *AtPDR11* confers improved paraquat resistance (Xi et al., 2012). Some L-type amino acid transporters also transport paraquat and confer paraquat resistance when their function is lost (Fujita et al., 2012; Li et al., 2013). Enhanced capability of ROS scavenging was also confirmed by Arabidopsis mutants (Chen et al., 2009; Luo et al., 2016). So far, other proposed mechanisms have not been confirmed.

The Detoxification Efflux Carriers (DTX)/Multidrug and Toxic Extrusion (MATE) subfamily genes are universally present in plants, animals and microorganisms and perform a variety of functions (Omote et al., 2006; Yuji Morita et al., 2000). There are more than 50 DTX members in the Arabidopsis genome (Li et al., 2002). The DTX subfamily proteins contain 3 to 12 signature transmembrane domains (Lu, 2016; Lu et al., 2018; Miyauchi et al., 2017; Tanaka et al., 2013), localize to various cell membranes or organelles (Dobritzsch et al., 2016; Li et al., 2002; Marinova et al., 2007; Serrano et al., 2013; Zhang et al., 2017; Zhang et al., 2014), and transport different sorts of substrates (Dobritzsch et al., 2016; Magalhaes et al., 2007; Roschzttardtz et al., 2011; Serrano et al., 2013). For example, AtDTX1 in plasma membrane participates in the efflux of plant-derived alkaloids, antibiotics and other toxic compounds in Arabidopsis (Li et al., 2002). DTX50 facilitates ABA efflux and thus modulates ABA sensitivity and drought tolerance (Zhang et al., 2014).

Recently, rice DG1, a MATE transporter too, was reported to play a key role in grain filling by mediating ABA long distance transport from leaf to caryopsis (Qin et al., 2021). DTX33 and DTX35 function as chloride channels and regulate cell turgor in Arabidopsis (Zhang et al., 2017). AtFRD3 and OsFRDL1 mediate citrate efflux and thus the response to iron deficiency (Durrett et al., 2007; Green and Rogers, 2004; Rogers and Guerinot, 2002; Yokosho et al., 2009). In addition, some DTXs have been shown to mediate various toxins resistance (Li et al., 2002; Omote et al., 2006). For instance, AtALF5 confers resistance to tetramethylammonium in yeast (Diener. et al., 2001). AtDTX1 enhances the tolerance to norfloxacin and heavy metal cadmium in *E. coli KAM3* strain (Li et al., 2002). Some DTXs/MATEs can positively regulate aluminum tolerance by enhancing citrate secretion (Doshi et al., 2017; Li et al., 2018; Li et al., 2017; Ma et al., 2018; Yokosho et al., 2011).

Here, we report the isolation and characterization of the gain-of-function mutant *paraquat tolerance 15-D* (*pqt15-D*) with enhanced resistance to paraquat. *PQT15* encodes AtDTX6 protein in the DTX/MATE family of Arabidopsis. A point mutation in *PQT15* results in G311E amino acid residue change, leading to paraquat resistance in *pqt15-D*. We further showed that overexpression of *DTX6/PQT15* also confered paraquat resistance in the wild type, whereas the *DTX6/PQT15* knockout mutants exhibited hypersensitive phenotype to paraquat. Heterologous expression of *DTX6* and *DTX6-D* in *E. coli* also significantly enhanced bacterial resistance to paraquat.

Paraquat efflux assays with isolated protoplasts and bacterial cells directly demonstrated the paraquat efflux activity of DTX6/PQT15. Taken together, our results demonstrate that DTX6 acts as a efflux transporter and confers paraquat resistance by exporting paraquat out of cytosol, therefore revealing a molecular mechanism of paraquat resistance that has not been reported before in higher plants.

## RESULTS

### Isolation and characterization of *pqt15-D* with enhanced paraquat resistance

To obtain more paraquat resistance loci, we carried out a near saturation mutagenesis with EMS (ethyl methyl sulfonate) in the Columbia ecotype wild type background (125,000 seeds mutagenized). Using our previously established method (Xi et al., 2012), we screened the whole M2 mutant library under 2 μM paraquat selection pressure and isolated dozens of paraquat resistant mutants. To identify novel loci of paraquat resistance, we performed genomic DNA sequencing of those known paraquat resistant loci for each mutant, including *PAR1* (Li et al., 2013), *RMV1* (Fujita et al., 2012), *AtPDR11* (Xi et al., 2012), and *PQT3* (Luo et al., 2016). To our surprise, the whole genetic screen ended up with only 2 major paraquat resistant loci. One is *par1* with more than 20 independent mutants each with a distinct mutation. The other is a novel locus represented by *pqt15-5* with 8 independent mutants but all with the same point mutation (Table S1).

We re-screened the progeny of the *pqt15-5* and confirm the paraquat resistance phenotype (Figure 1a). Then the mutant was backcrossed with Col-0 wild type to get BC1F1generation, and F1 plants were selfed to get BC1F2 generation. BC1F2 population was phenotyped for paraquat resistance on MS medium containing 2 μM paraquat (Figure 1b). The segregation ratio of resistance: sensitive ≈ 3:1 (survived: dead = 230:88; χ^2^ = 1.07< χ^2^ _(0.05)_ =3.84). The genetic analysis showed that the *pqt15-5* is caused by a single dominant nuclear gene mutation. Therefore, we renamed the mutant *pqt15-D*.

**Figure 1.**
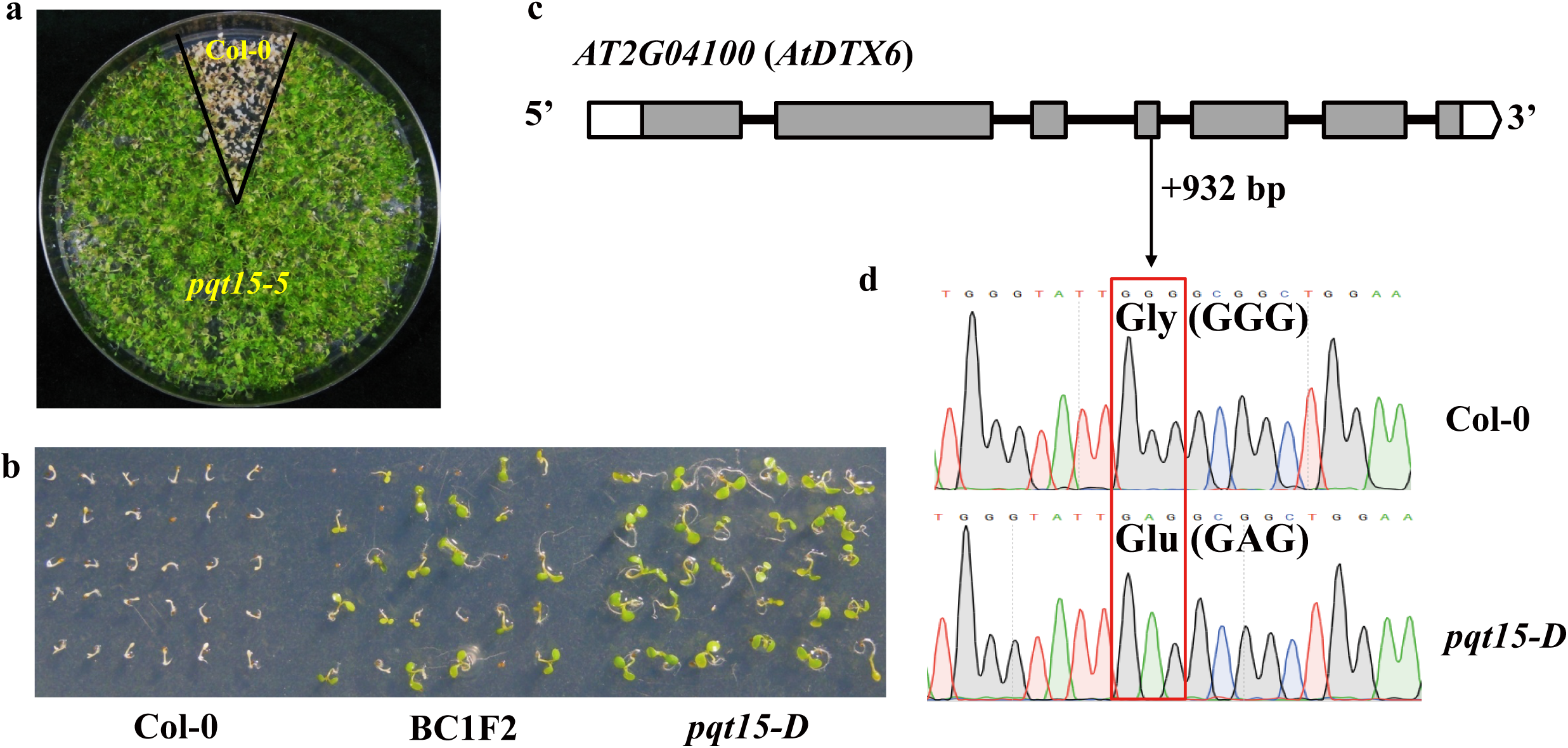
*pqt15-D* mutant and mutation identification. a. Paraquat resistance phenotype of *pqt15-D* mutant. Wild type (Col-0) and *pqt15-D* seeds were germinated and grown on MS medium containing 2 μM paraquat for 2 weeks before the image was recorded. b. Genetic analysis of the *pqt15-D* mutant. Segregation of paraquat resistance in BC1F2 population. Wild type (Col-0) was used as a negative control and *pqt15-D* as a positive control. c. Gene structure of the *AtDTX6*. White and gray boxes represent untranslated regions (UTRs) and exons, respectively, while black lines denote introns. d. Identification of the mutation in *AT2G04100/AtDTX6* of *pqt15-D* by sequencing. A red box indicates the codon with G-to-A point mutation. All the other 7 mutants (*pqt2-2, pqt8-4, pqt28-1, pqt63-1, pqt66-1, pqt72-1* and *pqt73-1*) have the same mutation in the same gene (Table S1).

To identify the mutation associated with paraquat resistance in the *pqt15-D* mutant, we pooled 50 paraquat resistant plants in BC1F2 population (more than 300 seeds) for whole-genome sequencing and identified a G-A substitution at position +932 bp in the coding region of *DTX6*, resulting in G311E replacement in amino acid sequence of DTX6 protein (Figure S1, Figure 1c, d). This same point mutation was identified in the other 7 independent mutants (Table S1) by genomic DNA sequencing, thus further confirming that this point mutation caused the paraquat resistance phenotype in *pqt15-D*.

### *DTX6/PQT15* is primarily expressed in roots and DTX6/PQT15 is mainly localized in plasma membrane

To examine the expression pattern of *DTX6/PQT15*, its transcript levels were detected for various plant tissues including roots, stem, rosette leaves, cauline leaves, flower, siliques, and seeds by quantitative RT-PCR analyses. The result shown that *DTX6/PQT15* was primarily expressed in the root and the expression levels were low in other tissues (Figure 2a).

**Figure 2.**
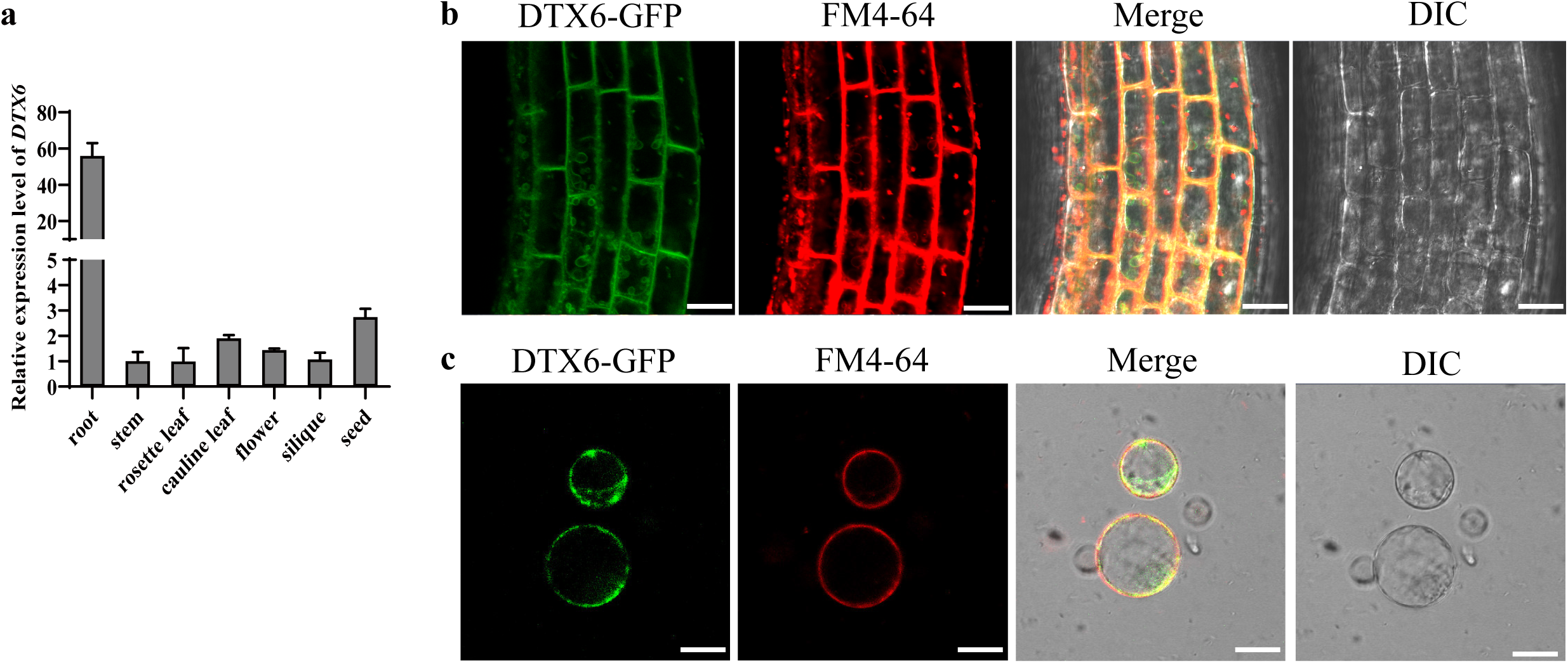
Expression pattern of *DTX6* and subcellular localization of DTX6. a. qRT-PCR analysis of *DTX6* expression levels in different tissues (root, stem, rosette leaves, cauline leaf, flower, silique, and seed). Values are mean ± SD (n= 3). b. Subcellular localization of *DTX6* in the root cells in the elongation zone of *DTX6pro:DTX6-GFP* transgenic plants. Bar =50 μm. c. Subcellular localization of DTX6 in the protoplasts prepared from the roots of *DTX6pro:DTX6-GFP* transgenic plants.

To analyze the subcellular localization of DTX6/PQT15, we generated *DTX6pro:DTX6-GFP* transgenic plants and found GFP signals in the plasma membrane and inner membranes of root cells (Figure 2b, c), indicating that DTX6-GFP fusion protein is localized in the plasma membrane and inner membranes in the root.

### DTX6/PQT15 confers paraquat resistance in Arabidopsis and *E. coli*

To verify genetically the function of DTX6, we obtained two knockout mutant lines (*dtx6-1* and *dtx6-2*) with CRISPR technology and three *35S-DTX6* transgenic Arabidopsis lines (OE-9, OE-13 and OE-17) in wild type background (Figure S2). These lines together with wild type control were subjected to paraquat resistance seed germination assay. There was no significant difference of survival among the lines when germinated on MS medium without paraquat. However, when germinated on MS medium containing paraquat of different concentrations (1.0, 1.5, and 2.0 µM), *dtx6-1* and *dtx6-2* lines were more sensitive to paraquat, whereas the OE-9, OE-13 and OE-17 showed significantly enhanced resistance to paraquat compared with wild type. Meanwhile, *pqt15-D* showed similar phenotype with OE lines (Figure 3a and b).

**Figure 3.**
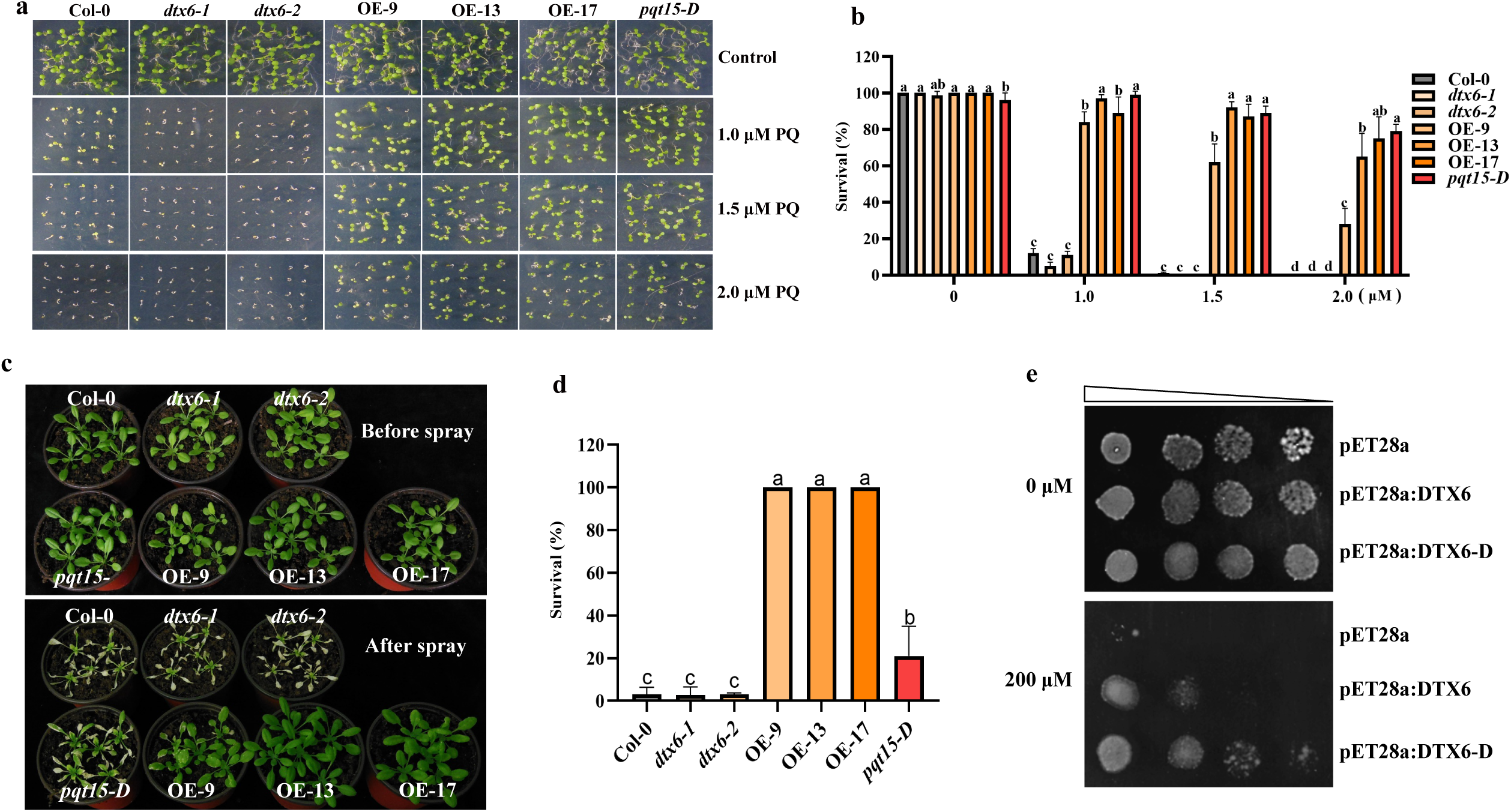
Enhanced DTX6 confers strong paraquat resistance. a. The phenotype of wild type (Col-0), *dtx6-1, dtx6-2*, OE-9, OE-13, OE-17, and *pqt15-D* germinated on MS medium containing different paraquat (PQ) concentration (0, 1.0, 1.5, and 2.0 μM) for 1 week. b. The survival ratio of wild type (Col-0), *dtx6-1, dtx6-2*, OE-9, OE-13, OE-17, and *pqt15-D*. Values are mean ± SD (n = 3, 25 seeds per replicate). Different letters indicate significant differences (P < 0.05; one-way ANOVA). c. Paraquat resistance assay. The phenotype of wild type (Col-0), *dtx6-1, dtx6-2*, OE-9, OE-13, OE-17 and *pqt15-D* grown in the soil for 4 weeks (top) and sprayed with 80 μM paraquat (bottom). Images were recorded 5 days after paraquat spray. d. The survival ratio of wild type (Col-0), *dtx6-1, dtx6-2*, OE-9, OE-13, OE-17 and *pqt15-D*. Values are mean ± SD (n= 3, 60 plants per replicate). Different letters indicate significant differences (P < 0.05; one-way ANOVA). e. Heterologous expression of *DTX6, DTX6-D* in *E. coli* confers paraquat resistance. *E. coli* Rosetta cells were transformed with the empty pET28a vector (negative control) or with pET28a containing the *DTX6* and *DTX6-D* cDNA, respectively, and grown on solid medium with 0 and 200 μM paraquat for 24 h before images were recorded.

We further assayed paraquat resistance of soil-grown plants. 4-week-old plants of *dtx6-1, dtx6-2*, wild type (Col-0), *pqt15-D*, and OE lines (OE-9, OE-13, and OE-17) grown in soil did not show significant morphological difference before paraquat spray (Figure 3c top). After paraquat spray, the knockout mutants (*dtx6-1* and *dtx6-2*) and wild type showed sensitive phenotype to paraquat, whereas the OE lines displayed significantly enhanced resistance with 100% vs. < 5% survival of wild type (Figure 3c bottom and d), but *pqt15-D* only showed slightly improved resistance to paraquat with 20% survival compared with wild type (Figure 3c and d) apparently due to the low expression level of *DTX6* in the leaf (Figure S3a & b, Figure 2a).

Moreover, *E. coli* expressing *DTX6* also displayed enhanced resistance to paraquat (200 µM) while *DTX6-D* confered even higher resistance (Figure 3e), further demonstrating the effect of the point mutation on paraquat resistance.

### DTX6/PQT15 effluxes paraquat out of the cell

Given that DTX6/PQT15 is a member of DTX/MATE subfamily transporters, which are known to participate in organic compounds and xenobiotic toxin efflux, we speculated that DTX6/PQT15 might function as a paraquat efflux transporter. To confirm the hypothesis, we conducted efflux assay with mesophyll protoplasts prepared from wild type (Col-0) and OE-13 plant leaves. Radiolabeled ^14^C-paraquat was preloaded into mesophyll protoplasts, and then ^14^C-paraquat was measured by monitoring the radioactivity retained in the protoplasts and released into the medium. The results showed that the rate of ^14^C-paraquat retention was significantly lower in OE-13 protoplasts (Figure 4a), while rate of ^14^C-paraquat release from OE-13 protoplasts was significantly faster than that from the wild type (Figure 4b).

**Figure 4.**
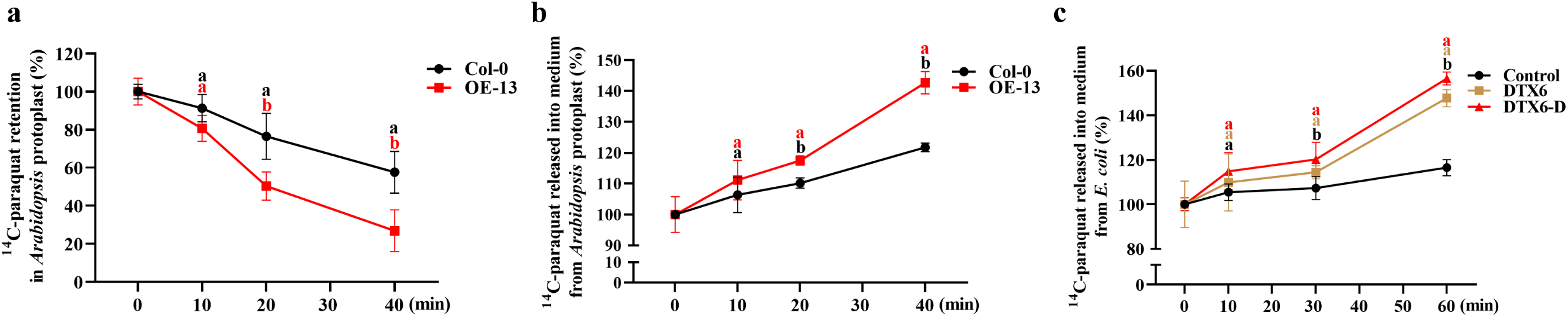
DTX6 effluxes paraquat out of the cell. a. Paraquat retention in the protoplasts. Paraquat efflux assay was conducted as described in Materials and Methods. Paraquat retained in the protoplasts was measured in Col-0 and OE-13 lines. Paraquat retention is defined as percent of the loaded 14C-paraquat in the protoplasts. Values are mean ± SD (n= 3). Different letters indicate significant differences (P < 0.05; one-way ANOVA). b. Paraquat release into the medium from the protoplasts. Paraquat efflux assay was conducted as described in Materials and Methods. Paraquat released into the buffer was measured in Col-0 and OE-13 lines. Paraquat release is defined as percent of the loaded 14C-paraquat in the protoplasts. Values are mean ± SD (n= 3). Different letters indicate significant differences (P < 0.05; one-way ANOVA). c. *E. coli* cells expressing *DTX6* and *DTX6-D* release more paraquat into medium than control (pET28a empty vector). Paraquat release is defined as percent of the loaded 14C-paraquat in bacterial cells. Values are mean ± SD (n= 3). Different

This was further confirmed by paraquat efflux assay with *E. coli* strains expressing *DTX6* or *DTX6-D*. Both DTX6 and DTX6-D were able to export paraquat out of the bacterial cells (Figure 4c).

### Increased negative charge on the surface of substrate entry domain may promote paraquat transport of DTX6-D

To understand the mechanism of DTX6-D enhanced paraquat transport, we simulated the 3-demension molecular models of DTX6 and DTX6-D based on the resolved crystal structure of AtDTX14 (Miyauchi et al., 2017), which shares a sequence identity of 50.9% with DTX6, and thus is considered as a suitable model for DTX6 and DTX6-D (Figure 5a). In the model, the residue Gly311 is localized in the entrance of the substrate tunnel. However, the variant G311E in DTX6-D would increase negative charge of the entrance (Figure 5b-d). Therefore, we hypothesize that the increased negative charge could favor the recognition of the positively charged paraquat and eventually increase the transport activity of DTX6. To prove this speculation, we mutated the Gly at 311 position to negatively charged glutamic acid, aspartic acid, and uncharged glutamine, respectively, then heterologously expressed in *E. coli*. Compared with the cells transformed with pET28a (empty vector), replacement with negatively charged glutamic acid or aspartic acid enhanced paraquat resistance of *E. coli*, whereas replacement with the uncharged glutamine did not enhance the resistance to paraquat compared with the wild type DTX6 on the medium with 200 μM paraquat (Figure 5e, f).

**Figure 5.**
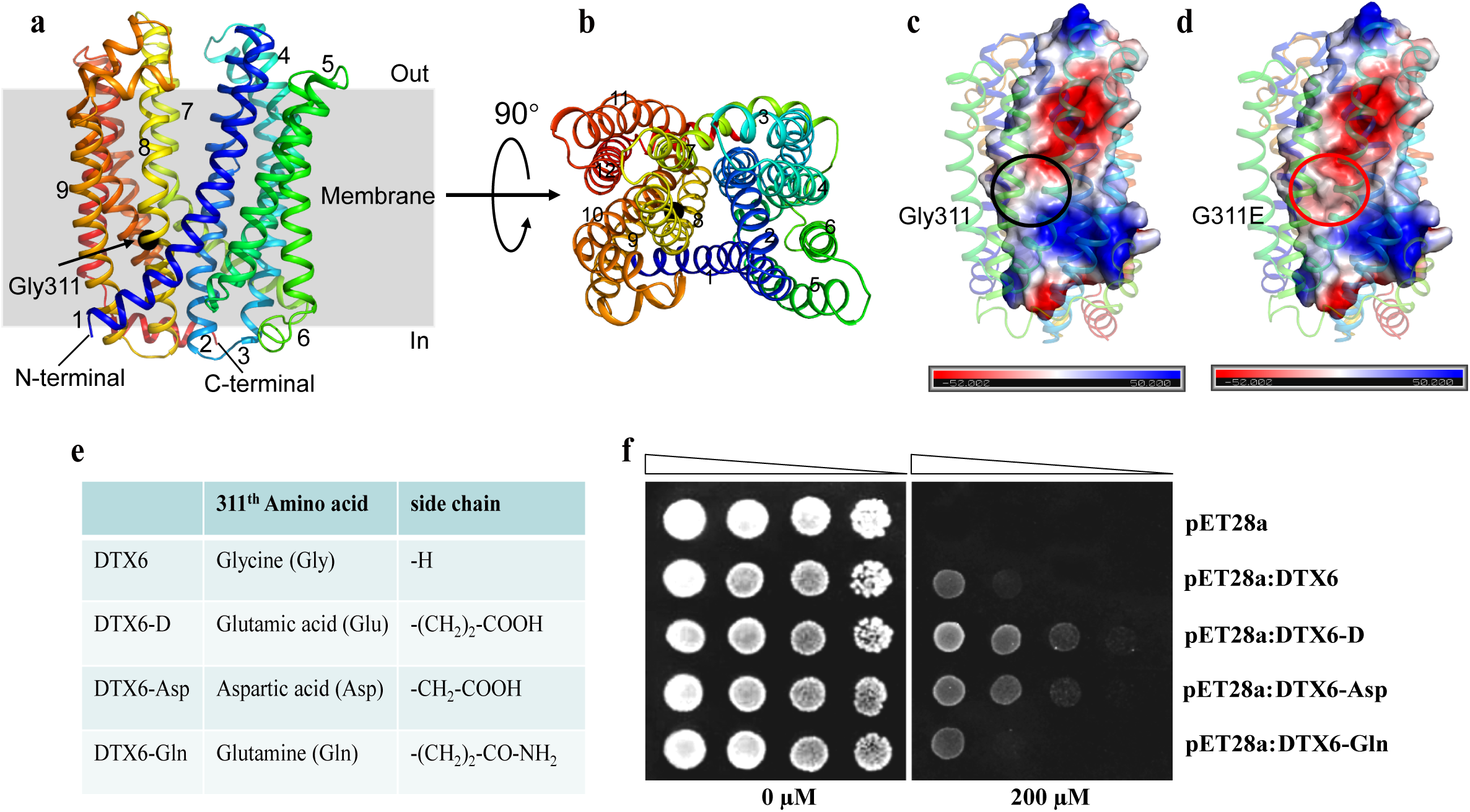
Increased negative charge on the surface of the substrate entry domain may promotes paraquat transport of DTX6-D. a. The structure model of DTX6 simulated by SWISS-MODEL from structure of AtDTX14 (Side view). The transmembrane helices are labeled and colored in rainbow. The residue Gly311 was indicated by black sphere. The red horizontal lines denote membrane with extracellular face (Out) and intracellular face (In). b. Top view of DTX6. c. Electrostatic surface potential map of DTX6. Gly311 is located in the entrance of transporter tunnel, the hydrophobic patch of Gly311 is highlighted by the black circle. The scale ranges from −50 kT/e (red) to 50 kT/e (blue). d. Electrostatic surface potential map of DTX6-D. Glu311 is located in the entrance of transporter tunnel, the negative charge patch of G311E is highlighted by red circle. The scale ranges from −50 kT/e (red) to 50 kT/e (blue). e. The 311th amino acid of DTX6 protein was mutated to Glu, Asp and Gln, respectively. f. *E. coli* Rosetta cells were transformed with the empty pET28a vector (negative control), or with pET28a containing the *DTX6* wild type cDNA or *DTX6-D, DTX6-Asp*, and *DTX6-Gln* cDNA with the mutation causing G311E, G311D, and G311N amino acid change respectively, and grown on solid medium with 0 and 200 μM paraquat for 24 h before images were recorded.

## DISSCUSSION

In this study, we report the isolation and characterization of a gain-of-function mutant *pqt15-D* (or *dtx6-D*) with enhanced paraquat resistance. Using the MutMap method (Takagi et al., 2015), we identified a G to A point mutation at position +932 in the coding region of *DTX6/PQT15* encoding a MATE transporter, which caused the enhanced paraquat resistance phenotype in *pqt15-D* (Figure 1, Figure S1). Moreover, the enhanced paraquat resistance can also be achieved by overexpressing *DTX6* in Arabidopsis and *E. coli* (Figure 3). We further revealed the molecular mechanism underlying the DTX6/PQT15-conferred paraquat resistance by paraquat efflux assays with mesophyll protoplasts and *E. coli* cells, which clearly showed that DTX6 and DTX6-D were able to export paraquat out of the cell (Figure 4), reducing cellular paraquat concentration and thus alleviating its toxic effects. Consistent with this, DTX6-GFP fusion protein was mainly localized to the plasma membrane (Figure 2). Therefore, our results have unraveled a molecular mechanism of paraquat resistance in higher plants that has not been reported before.

Considering DTX6 as a member of MATE transporter subfamily, which has been reported to transport kinds of organic compounds in bacteria, plants and animals (Kuroda and Tsuchiya, 2009; Li et al., 2002; Omote et al., 2006), DTX6 may be capable of effluxing paraquat from the cytosol to the extracellular space. Indeed, as shown by the efflux assays, the wild type DTX6 is able to export paraquat out of the protoplasts (Figure 2a, b), but the paraquat efflux activity is likely too low to confer paraquat resistance in wild type. Once its activity is enhanced either by the point mutation as in the *pqt15-D* mutant or elevated expression as in the overexpression lines, strong paraquat resistance can be conferred (Figure 3). What are the natural substrates of DTX6 is an interesting question to pursue. It is conceivable that its natural substrates are quite likely positively charged compounds.

The subcellular localization of DTX6 is in agreement with its role as an efflux transporter. Our *DTX6pro:DTX6-GFP* transgenic plants clearly show that TDX6-GFP fusion protein is mainly in the plasma membrane in the root. However, GFP signals also appeared in the inner membranes, indicating the possibility of other subcellular location of DTX6. Since the *DTX6* promoter activity is very low in the leaf, we could not detect GFP signals in the leaf with our *DTX6pro:DTX6-GFP* transgenic plants.

Considering the strong paraquat resistance of the OE lines, it is conceivable that DTX6 would similarly localize in plasma membrane in the leaf but would be interesting to see where else DTX6-GFP is subcellularly localized.

How could a single amino acid change (G311E) of DTX6 afford Arabidopsis resistant to paraquat? Since the G311E single amino acid replacement predicted in the substrate entry domain is solely responsible for enhanced paraquat resistance in the mutant, we reasoned that electrical charge from neutral to negative would favor the binding of DTX6 to paraquat, thus increase DTX6 protein efflux activity and confer higher resistance to paraquat (Figure 3e). Indeed, G311D substitution in DTX6 could also confer enhanced paraquat resistance (Figure 5f), suggesting that a more negatively charged surface at the substrate entry would favor paraquat interaction with DTX6, which may enhance efflux activity of DTX6.

In summary, with the *pqt15-D* mutant, we unraveled DTX6 as a paraquat efflux transporter. No paraquat efflux transporters of higher plants have been reported before. Paraquat efflux out of the cell represents a new mechanism for paraquat resistance in higher plants. We have demonstrated that enhanced DTX6/PQT15 via either the G311E amino acid alteration as in *pqt15-D* or elevated expression of *DTX6/PQT15* as in the OE lines would confer strong resistance to paraquat in Arabidopsis and *E. coli*. Undoubtedly, this will provide a promising candidate gene for engineering paraquat resistance crops.

## MATERIALS AND METHODS

### Plant materials and growth conditions

Arabidopsis Columbia ecotype (Col-0) was used in this study. Two mutants *dtx6-1* and *dtx6-2* were obtained via CRISPR/Cas9 technology (Figure S2a-c). The transgenic plants were obtained by transferring *DTX6pro:DTX6-GFP* and *35S-DTX6* constructs into Arabidopsis using the floral dip method (Bent., 2000). Plants were grown in soil under greenhouse condition (22°C; 16/8 h light/dark cycle for long-day; 8/16 h light/dark for short-day; with light intensity 100 μM m^-2^ s^-1^) or MS medium containing 0.5% agar and 1% sucrose in the growth chamber (22 °C; 16/8 h light/dark cycle for long-day; 100 μM m^-2^ s^-1^ light intensity).

### Genetic screen and identification of mutation associated with paraquat resistance in the mutants

More than 1,000,000 EMS-mutagenized M2 seeds were germinated and grown for 7-10 days on MS medium containing 2 µM paraquat to screen for paraquat resistant mutants. In the germination phase, the vast majority of seedlings are bleached or cannot product roots, whereas candidate paraquat resistant mutants stay green and produce roots. The candidate paraquat resistant mutants were transferred to soil for M3 seeds. M3 seeds were rescreened under same condition, and then were used to backcross with wild type for F1 seeds and F2 was further obtained through F1 selfing.

The F2 population was phenotyped by using the same screening condition and 50 resistant plants were pooled to extract genomic DNA for whole genome re-sequencing by the BSA method (Abe et al., 2012; Takagi et al., 2015). Genome re-sequencing was commercially done by Weifen (Anhui) Gene Technology Co. (Hefei, China).

### Paraquat resistance assay of seed germination

For paraquat resistance assay of seeds germination, seeds were surface-sterilized for 10-15 min in 10% bleach, washed five times with sterile water, and seeds were vernalized at 4°C for 48 h. After vernalization, seeds were plated on MS solid medium with 1% (w/v) sucrose, 0.5% (w/v) agar or MS medium containing different concentration paraquat (1.0, 1.5 and 2.0 µM). After one week, the survived was counted. Survival was defined as percent of seedlings with green cotyledon.

### Paraquat spray assay of soil grown plants

4-weeks-old plants grown in soil were sprayed with 80 µM paraquat solution. After 5 days, the survived was counted. Survival was defined as percent of plants with green leaves.

### Quantitative RT-PCR analyses

Total RNA was extracted from different tissues of plants with Trizol method (Invitrogen, Carlsbad, USA). After that total RNA is reversely transcribed into cDNA by TransScript Kit (TaKaRa, Dalian, China). The specific primers were used for RT/qRT-PCR (Table S1).

### Subcellular localization of DTX6-GFP fusion protein

Roots of *DTX6pro:DTX6-GFP* transgenic plants were observed by using ZEISS880 confocal laser scanning microscope as described (Chaiwanon et al., 2016; Zhao et al., 2020). Protoplasts were prepared from the transgenic roots as described (Yoo et al., 2007) and observed as above.

### ^14^C-paraquat efflux assay in Arabidopsis protoplasts

Arabidopsis mesophyll protoplasts were isolated from Arabidopsis rosette leaves of Col-0 and OE-13 lines grown for 4 weeks in a short day photoperiod as described (Yoo et al., 2007). About 6×10^6^ of mesophyll protoplasts were incubated for 1 h in 2 ml bathing W5 solution (154 mM NaCl, 125 mM CaCl_2_, 5 mM KCl and 2 mM MES, pH 5.7) with 3 µM ^14^C-paraquat (American Radiolabeled Chemicals, Inc., Saint Louis, USA) at room temperature. Then, the mesophyll protoplasts were washed three times with ice cold bath W5 solution to remove unloaded ^14^C-paraquat and resuspended in 13 ml bath W5 solution with protoplast concentration of 5×10^5^ per ml. ^14^C-paraquat release from the protoplasts was monitored by removing 1 ml aliquot sample in triplicate at different time points (0, 10, 20 and 40 min) at room temperature.

Immediately, the aliquoted samples were pulse spinned to separate protoplasts from the bath solution, then the protoplast pellet and supernatant were separately mixed with 5 ml scintillation solution to measure radioactivity by liquid scintillation counter (Perkin-Elmer Tri-carb 2910TR, Waltham, USA).

### Paraquat resistance and ^14^C-paraquat efflux assays of *E. coli* ROSETTA strain expressing DTX6 and DTX6-D

The *DTX6, DTX6-D, DTX6-Asp*, and *DTX6-Gln* cDNA were constructed into the pET28a vector, respectively, and introduced into the *E. coli* ROSETTA strain and grown on solid LB medium with 50 mg/ml kanamycin antibiotics, single clones were picked, incubated in liquid medium containing 50 mg/ml kanamycin and 1 mM IPTG to OD_600_ = 0.4, then 2 µl of the 1:10 diluted culture was spotted on LB solid medium containing 50 mg/ml kanamycin, 1 mg/ml IPTG, and different concentrations of paraquat and incubated for 24 h at 37°C before colony growth was checked. The ROSETTA strain with pET28a empty plasmid was used as a control.

A ^14^C-paraquat transport assay was performed as described previously (Zhang et al., 2014). DTX6 and DTX6-D were expressed in *E. coli* ROSETTA strain, respectively, and then 5 µM ^14^C-paraquat was loaded into the ROSETTA strains with pET28a (control), pET28a-DTX6, and pET28a-DTX6-D plasmid, respectively.

Paraquat release from bacterial cells was monitored by removing aliquot samples at different time points (0, 10, 30 and 60 min) at room temperature. After that, the sample was pulse spinned, then the supernatant was mixed with 5 ml scintillation solution for monitoring radioactivity by liquid scintillation counter (Perkin-Elmer Tri-carb 2910TR, Waltham, USA).

### 3D-molecular models of DTX6 and DTX6-D

Using AtDTX6 homologous proteins AtDTX14 as the most suitable molecular structural templates (Protein Data Bank accession 5Y50). AtDTX14 has norfloxacin export activity like AtDTX1 (Li et al., 2002; Miyauchi et al., 2017). 3D-molecular models of DTX6 and DTX6-D were constructed with Phyre2 software.

### Statistical analysis

Statistical significance was evaluated at the 0.05 probability level using Student’s t-test.

## AUTHOR’S CONTRIBUTION

JQX and CBX designed the experiments. JQX performed experiments and data analysis, and wrote the manuscript. TN, AJ, MA, QQL, ZYZ, ZSZ and JW contributed to assist in completing part of the experiment. LW, YXC, ZSY, and LFS contributed to construct 3D-molecular models. CBX supervised the project and revised the manuscript.

## ACKNOWLEDGEMENTS

This work was supported by grants from National Natural Science Foundation of China (grant no. 31770273). The authors thank Dr. Xiaochun Ge (Fudan University, Shanghai, China) for her assistance with microscopic observation of DTX6-GFP localization and helpful discussions.

## Supplemental information

Figure S1. The mapping of the mutation responsible for *pqt15-D* phenotype based on MutMap analysis.

Figure S2. Identification of *dtx6* mutants.

Figure 3S. Relative expression level of *DTX6*/*DTX6-D* gene in leaves of 1-week-old and 4-weeks-old plants, respectively.

Table S1. Alleles of 2 major paraquat resistance loci in Arabidopsis

Table S2. Primers sequences used in this study.

## Notes

### Competing Interest Statement

The authors have declared no competing interest.

