## supplemental info for "Enhanced MATE transporter DTX6/PQT15 confers paraquat resistance"

**Supplemental information for “Enhanced MATE transporter DTX6/PQT15 confers paraquat resistance” by Xia et al.**

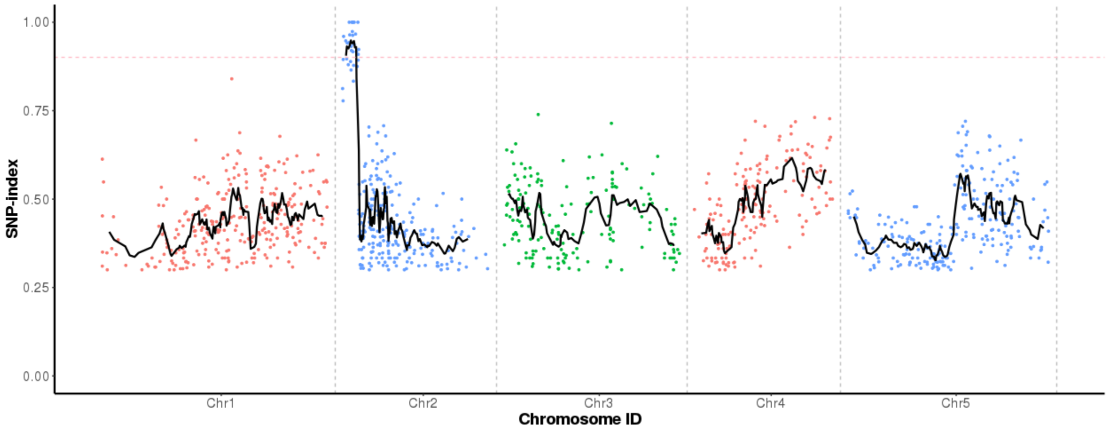


**Figure S1. The mapping of the mutation responsible for *pqt15-D* phenotype based on MutMap analysis.**

SNP-index plot of the *Arabidopsis thaliana* whole-genomic by MutMap analysis, showing a genomic region with the highest SNP-index peak harboring the candidate mutation. DNA bulked from 50 paraquat tolerant F2 progeny obtained from the backcross with Col-0 and *pqt15-5* was used for sequencing and MutMap analysis. The black line represents the sliding window average values of SNP indices of 1-Mb intervals with a 10 kb increment. The pink dotted line is the index (0.9) threshold line.

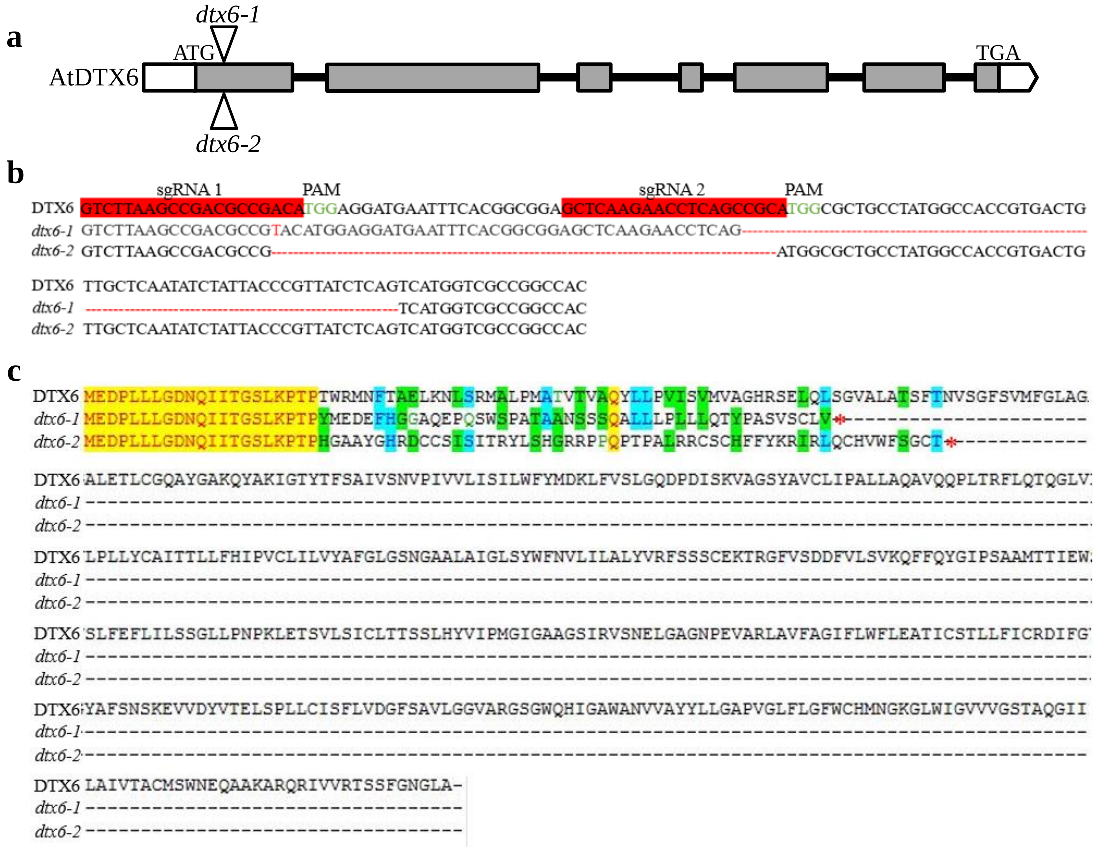


**Figure S2. Identification of *dtx6* mutants.**

a. Schematic representation of *DTX6* gene and location of CRISPR-Cas9 edited sites indicated by triangles.

b. Sequence around two CRISPR/Cas9 target sites ( sgRNA1 and sgRNA2 ), and the mutated nucleotide sequences of *dtx6-1* and *dtx6-2* lines.

c. The protein sequences translated from the mutated nucleotide sequences of *dtx6-1* and *dtx6-2* lines. The red stars represent stop codon.

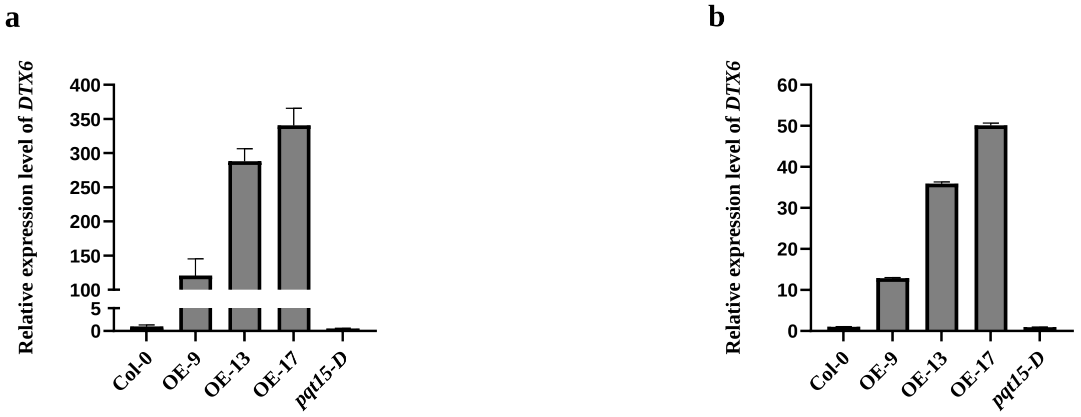


**Figure 3S. Relative expression level of *DTX6*/*DTX6-D* gene in leaves of 1-week-old and 4-weeks-old plants, respectively.**

a. The *DTX6*/*DTX6-D* gene expression level in leaves of 1-week-old plants.

b. The *DTX6*/*DTX6-D* gene expression level in leaves of 4-weeks-old plants.

**Table S1. Alleles of 2 major paraquat resistance loci in Arabidopsis**

| gene | Mutant lines | Mutation site | Base before mutation | Base after mutation | Amino acid before mutation | Amino acid after mutation |
|---|---|---|---|---|---|---|
| AT1G31830 | *pqt1-2* | +593 bp | TCC | TTC | Ser | Phe |
| | *pqt1-4* | +731 bp | TCG | TTG | Ser | Leu |
| | *pqt2-3* | +349 bp | GTT | ATT | Val | Ile |
| | *pqt2-4* | +684 bp | TGG | TGA | Trp | Stop |
| | *pqt4-18* | +375 bp | TGG | TGA | Trp | Stop |
| | *pqt5-1* | +310 bp | GGA | AGA | Gly | Arg |
| | *pqt6-1* | +726 bp | TGG | TGA | Trp | Stop |
| | *pqt6-2* | +203 bp | GGT | GAT | Gly | Asp |
| | *pqt8-1* | +1019 bp | GCT | GTT | Ale | Val |
| | *pqt15-2* | +727 bp | GAC | AAC | Asp | Asn |
| | *pqt16-2* | +325 bp | GAG | AAG | Glu | Lys |
| | *pqt21-1* | +302 bp | GCG | GTG | Ala | Val |
| | *pqt35-1* | +799 bp | GTG | ATG | Val | Met |
| | *pqt35-2* | +737 bp | AGT | AAT | Ser | Asn |
| | *pqt58-1* | +875 bp | TGG | TAG | Trp | Stop |
| | *pqt58-2* | +393 bp | TGG | TGA | Trp | Stop |
| | *pqt61-3* | +1081 bp | GGG | AGG | Gly | Arg |
| | *pqt65-2* | +1135 bp | GAG | AAG | Glu | Lys |
| | *pqt73-3* | +332 bp | GGT | GAT | Gly | Asp |
| | *pqt73-4* | +304 bp | GAG | AAG | Glu | Lys |
| | *pqt76-1* | +935 bp | TGG | TAG | Trp | Stop |
| | *pqt77-1* | +711 bp | TGG | TGA | Trp | Stop |
| | *pqt78-6* | +410 bp | GGT | GAT | Gly | Asp |
| AT2G04100 | *pqt2-2* | +932 bp | GGG | GAG | Gly | Glu |
| | *pqt8-4* | +932 bp | GGG | GAG | Gly | Glu |
| | *pqt15-5* | +932 bp | GGG | GAG | Gly | Glu |
| | *pqt28-1* | +932 bp | GGG | GAG | Gly | Glu |
| | *pqt63-1* | +932 bp | GGG | GAG | Gly | Glu |
| | *pqt66-1* | +932 bp | GGG | GAG | Gly | Glu |
| | *pqt72-1* | +932 bp | GGG | GAG | Gly | Glu |
| | *pqt73-1* | +932 bp | GGG | GAG | Gly | Glu |

### Table S2. Primers sequences used in this study.

| Name | Primer Sequence | vector |
|---|---|---|
| CRISPR/Cas-DTX6-Guide 1 | P1: ATTG GTCTTAAGCCGACGCCGACA | pCAMBIA1300-pYAO: Cas9 (for DTX6 gene editing) |
| | P2: AAAC TGTCGGCGTCGGCTTAAGAC | |
| CRISPR/Cas-DTX6-Guide 2 | P1: ATTG GCTCAAGAACCTCAGCCGCA | |
| | P2: AAAC TGCGGCTGAGGTTCTTGAGC | |
| HA-35S-DTX6 | P1: GGGGACAAGTTTGTACAAAAAAGCAGGCTATGG AAGATCCACTTTTATTGGG | pCB2004 (for DTX6 gene Overexpression) |
| | P2: GGGGACCACTTTGTACAAGAAAGCTGGGT TCAAGCAAGTCCATTGCCAAATG | |
| DTX6pro-DTX6-GFP | P1: GGGGACAAGTTTGTACAAAAAAGCAGGCTTCTG AAGAAAGGAAATAACAAAGC | pMDC110 (for *DTX6pro*:DTX6-GFP) |
| | P2: GGGGACCACTTTGTACAAGAAAGCTGGGTAGCAA GTCCATTGCCAAATGAA | |
| 35S-DTX6-GFP | P1: GGGGACAAGTTTGTACAAAAAAGCAGGCTATGG AAGATCCACTTTTATTGGG | pGWB5 (for 35S:DTX6-GFP) |
| | P2: GGGGACCACTTTGTACAAGAAAGCTGGGTCAGC AAGTCCATTGCCAAATGAA | |
| pET28a-DTX6-BamH1 pET28a-DTX6-Not1 | P1: CGGGATCCATGGAAGATCCACTTTTATTGGG | pET28a (for DTXs protein expression) |
| | P2: ATAAGAATGCGGCCGCTCAAGCAAGTCCATTGCCAAATG | |
| pet28a-at2g04090-BamH1 pet28a-at2g04090-Sal1 | P1: CGGGATCC ATGGAAGATCCACTTTTATTGGG | |
| | P2: ACGCGTCGAC TCACTCCAATGTTCTTCCAACTA | |
| pET28a-DTX6-Gln | P1: ATTCCAATGGGTATTCAGGCGGCTGGAA | |
| | P2: GAATACCCATTGGAATGACATAGTGGAGA | |
| pET28a-DTX6-Asp | P1: TTCCAATGGGTATTGATGCGGCTGGAAGTA | |
| | P2: ATCAATACCCATTGGAATGACATAGTGGA | |
| DTX6-q-PCR | P1: GACATGGAGGATGAATTTCACG | |
| | P2: CCGATTTTCGCGTATTGTTTTG | |
| ACTIN2 | P1: TGAAGTATCCTATTGAGC | |
| | P2: CTTTGGGTTAAGAGGAGCCTCG | |
| β—tubulin8 | P1: CTTAAGCTCACCACTCCAAGCT | |
| | P2: GCACTTCCACTTCGTCTTCTTC | |
